# Sensory substitution reveals a manipulation bias

**DOI:** 10.1101/816835

**Authors:** AT Zai, S Cavé-Lopez, M Rolland, N Giret, RHR Hahnloser

## Abstract

Sensory substitution is a promising therapeutic approach for replacing a missing or diseased sensory organ by translating inaccessible information into another sensory modality. What aspects of substitution are important such that subjects accept an artificial sense and that it benefits their voluntary action repertoire? To obtain an evolutionary perspective on affective valence implied in sensory substitution, we introduce an animal model of deaf songbirds. As a substitute of auditory feedback, we provide binary visual feedback. Deaf birds respond appetitively to song-contingent visual stimuli, they skillfully adapt their songs to increase the rate of visual stimuli, showing that auditory feedback is not required for making targeted changes to a vocal repertoire. We find that visually instructed song learning is basal-ganglia dependent. Because hearing birds respond aversively to the same visual stimuli, sensory substitution reveals a bias for actions that elicit feedback to meet animals’ manipulation drive, which has implications beyond rehabilitation.

## Main Text

Sensory substitution is a transformation of stimuli from one sensory modality into another one^1^. Such transformation can be used as a therapeutic approach towards restoring perception from a defective sensory modality^2^. This approach has gained much interest in recent years thanks to both advances in technology and the remarkable cross-modal flexibility of the central nervous system^3–5^. However, one of the main obstacles hindering the wide adoption of substitution devices has been the amount of training necessary to make use of the new sensory input; in fact, blind subjects often give up using a substitution device before reaching a reasonable proficiency level because they feel overwhelmed and frustrated^4^.

How can this situation be remedied, and which are the general design principles that need to be respected for sensory substitution to be willingly adopted? Currently, the motivational consequences inherent in sensory substitution are poorly understood, partly because we are lacking a theory that would predict how a subject will respond to substituting input. One key question is whether substitution will increase or decreases the affective valence of a given motor action^6,7^. Ideally, we would like to know beforehand about actions that will suffer from a decrease in valence and therefore will be avoided by subjects. Vice versa, if we could predict the actions that will experience a boost in valence from substitution, we could provide better treatments to support skilled behaviors such as speech in the deaf.

This question concerns the motivational system that is best served by substitution. One idea is that sensorially deprived subjects desire highly informative feedback about their actions. For example, substituting input could help subjects to reduce uncertainties inherent in their motor output and allow them to make better action choices. Accordingly, the artificial sensory input should perfectly differentiate among distinct action outcomes. Formally, substitution may elicit the desire to explore^8–10^, which is to seek knowledge about actions’ effects. According to this knowledge seeking view, subjects will preferentially choose actions with uncertain outcomes^11^ or high predicted information gain^12–14^.

Another idea is that adaptive responses to substitution may focus on the intrinsic goal of manipulating the environment^15^ rather than to obtain knowledge. For example, subjects may be drawn towards actions for the sole reason that the latter trigger a significant sensory input. Substitution could thus reveal a drive for impact^16^, which is to preferentially choose actions with noticeable effect.

To test whether knowledge-seeking or impact-seeking better explains adaptive responses to sensory substitution, in songbirds we partially replace auditory feedback from a complex vocal behavior by visual feedback. We modified a widely applied operant conditioning paradigm involving the pitch of a song syllable. Instead of using short white-noise bursts played through a loudspeaker^17,18^, we substitute auditory by visual feedback by briefly switching off the light in the sound-isolation chamber of the singing bird whenever the pitch of a targeted syllable was below (or above) a threshold, Fig. 1. We set the pitch threshold for light-off every morning to the median pitch value on the previous day. We investigated whether adult male zebra finches deafened using bilateral cochlea removal would show any kind of reaction in response to such pitch substitution by light-off (LO). We evaluated bird’s responses to substitution in terms of d’ values, which are average daily pitch changes normalized by their standard deviations (see Methods). From these values, we inferred the affective valence of substituted feedback: whether it is neutral, aversive, or appetitive.

**Fig. 1:**
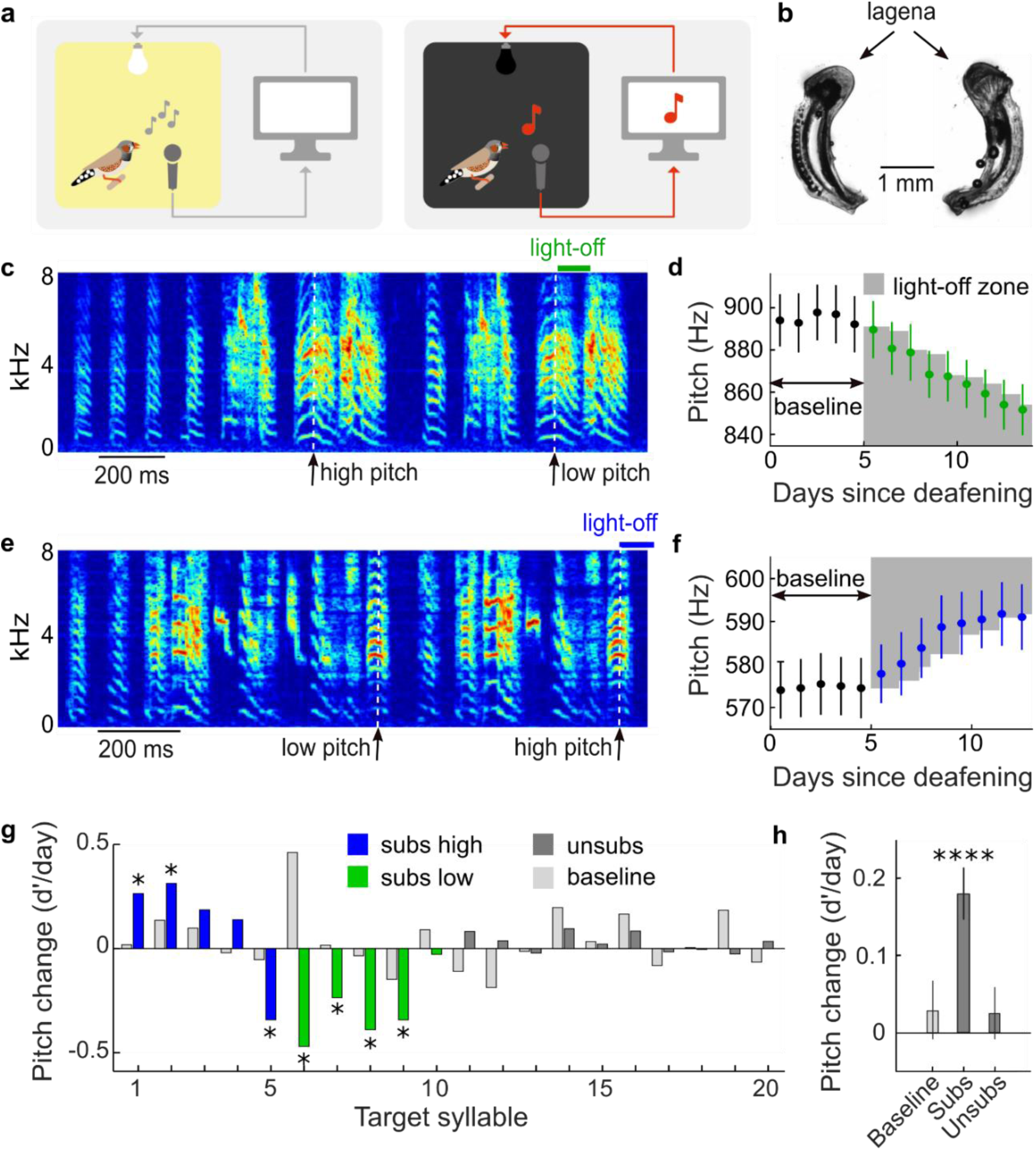
Light-off stimuli are positive reinforcers of vocal pitch in deaf songbirds. **a**, Schematic of the experiment. A singing deaf bird inside a sound isolation chamber (left) experiences light off for a duration in the range of 100-500 ms (right) when the pitch of one of its song syllables (red note) exceeds a given threshold. **b**, Example picture of a pair of surgically removed cochlea. Complete deafness was confirmed by presence of the osseous spiral lamina and by verification of an intact loop including the lagena. **c, e**, Example song spectrograms in birds b2y2 (**c**) and b2p19 (**e**) with substituted feedback for low-pitched (**c**) and high-pitched (**e**) syllable renditions (the time points of pitch measurement are indicated by white dashed lines). **d, f**, Daily mean pitches (dots) and their standard deviations during baseline (black) and during substitution (green: low-pitch subs, **d**; blue: high-pitch subs, **f**). The light-off pitch zones are indicated by gray shading. The birds adapted the pitch in the direction of increasing light-off rate. **g**, Histograms of average daily pitch changes during substitution in birds with high-pitch substitution (subs high, blue, n=5 birds, the first bar corresponds to b2p19 shown in **e** and **f**), low-pitch substitution (subs low, green, n=5 birds, the 8^th^ bar corresponds to b2y2 shown in **c** and **d**), and in control birds without substitution (unsubs, dark grey, n=7 birds, target syllables 11 and 14 stemmed from the same bird and so did syllables 13+18 and 15+19). The light grey bars to the left of the colored bars indicate the average daily pitch change for each syllable during the last 5 baseline days. The asterisks indicate subs birds with significant pitch changes compared to controls (two sample, two-sided t-test, p<0.05). **h**, Shown are the three fixed-effect terms of a mixed linear effect model and their standard errors (see main text for details). The bars indicate the daily change in pitch (d’/day) during baseline (left, p=0.44), during substitution in the direction of increasing light-off rate (subs, p=2.0*10^−7^), and in control birds (unsubs, right, p=0.46).

### Substituted feedback appetitively reinforces vocal pitch

Because deafening by itself may induce a slow pitch drift with a nonzero bias^19,20^, we evaluated pitch responses in comparison to unsubstituted deaf control (unsubs) birds. Pitch changes in 7/10 subs birds significantly deviated from the changes in control animals in matched time periods (p<0.05 in 7 of 10 subs birds, two sample, two-tailed t-test of pitch change per day, see Methods, Fig. 1g).

Interestingly, subs birds tended to be attracted by light-off, because all birds except one changed their pitch in the direction of increasing LO rate, Fig. 1g. If the direction of pitch drift were random in each bird with probability ½ in each direction (binomial model), then 9 of 10 birds would drift in the same direction in less than 1% of cases, corresponding to a p-value smaller than 0.01, whence the attraction was a non-random effect.

To account for individual variability, we fitted mixed linear effect models to the pitch data. The models contained three fixed terms: one term for the early time period before substitution and one term each for the late time periods in subs and unsubs birds. In addition, there was one random term for each syllable (n=17 birds, 20 target syllables, see Methods). We found that relative to baseline (early period), subs birds exhibited pitch changes of 0.18 d’/day in the direction of increasing LO rate (nonzero fixed effect, p=2.0*10^−7^, SE=0.03, tstat=5.33, DF=282, Fig. 1h), whereas unsubs birds did not change pitch (0.02 d’/day, p=0.46, SE = 0.03, tstat=0.74, DF=282).

Syllables in deaf birds remained stable over the short period of the experiment; changes were specific to pitch but did not affect other sound features (p>0.05, two-tailed t-test, duration, frequency modulation, amplitude modulation, and entropy, see Methods), see Supplementary Fig. S1. In combination, these results indicate that in deaf birds, substituted feedback is an appetitive reinforcer of song.

### The same stimulus aversively reinforces vocal pitch in hearing birds

We also evaluated adaptive pitch responses in hearing birds. A small number of hearing birds responded to light-off: Pitch changes in two of 12 birds significantly exceeded the spontaneous pitch drift in hearing controls (no LO, p<0.05, two-tailed t-test on pitch changes per day, see Methods, Fig. 2c). A mixed linear effect model revealed that hearing birds significantly changed pitch in the direction of decreasing LO rate (−0.08 d’/day in the direction of subs, nonzero fixed effect of LO, p=1.6*10^−4^, DF=385, SE=0.02, tstat=-3.81, N=24 syllables from 24 birds including 12 controls), implying that overall, light-off was aversive in hearing birds. In combination, our findings demonstrate that deafening causes an inversion of affective valence of LO reinforcers.

**Fig. 2:**
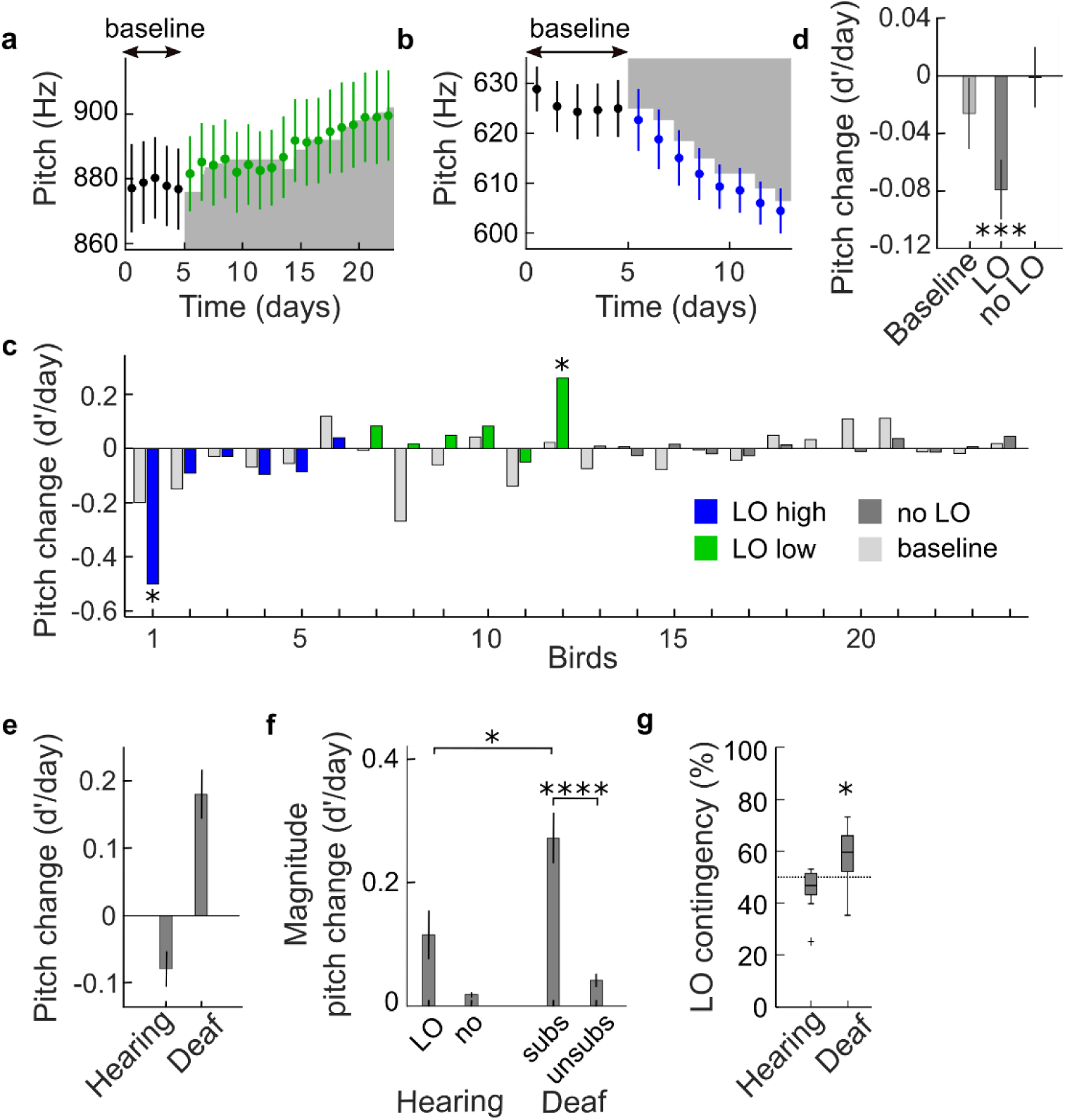
In hearing birds, the valence of light-off reinforcers is negative. **a, b**, Hearing birds change pitch in the direction of decreasing LO rate, here shown for a low-pitch light-off (LO low) bird (bird b2y2, **a**) and a LO high bird (bird p6s6, **b**). Legend as in Fig. 1d, f. **c**, Histograms of average daily pitch changes in LO high (blue, n=6), LO low (green, n=6), and in no LO birds (gray, n=12). **d**, The three mixed linear fixed effect terms and their standard errors. The bars indicate the daily change in pitch (d’/day) during baseline (left, p=0.28), during LO exposure in the direction of increasing LO rate (middle, p=1.6*10^−4^), and in control birds (no LO, right, p=0.96). **e**, Average directed pitch changes in LO (hearing) vs subs (deaf) birds. Hearing birds changed their pitch away from the LO zone (decreasing the number of renditions with LO) and deaf birds towards the LO zone (increasing the number of renditions with LO). The error bars indicate the standard errors of the mean. **f**, The magnitude of average pitch change is larger in subs birds than in LO birds (* indicates p<0.05 for average magnitude, two sample two-sided t-test), and much larger than in unsubs birds (in time-matched periods, **** indicates p<0.001). The error bars indicate standard errors of the mean. **g**, Distribution of average daily LO contingency for hearing and deaf birds. Deaf birds triggered light-off in more than 50% of cases (58%, p=0.04, tstat=2.45, SD=0.11 DF=9, two-tailed t-test) whereas in hearing birds, the LO contingency did not significantly deviate from 50% (46%, p=0.08, tstat=-1.91, SD=0.08, DF=11, two-tailed t-test).

Valence inversion was also signaled by the contrasting effects of light-off on singing rates. Subs birds produced 280% or 1164 more song motifs on the last three days of substitution than their deaf controls (non-zero fixed effect of light-off, p=5*10^−4^, DF=58, SE=318, tstat=3.66, N=20 syllables including 10 syllables from controls), suggesting that substitution increases the motivation of deaf birds to sing. Hearing (LO) birds, by contrast, had a tendency to sing 10% or 201 fewer song motifs per day than their hearing controls (fixed effect of light-off, p=0.72, DF=70, SE=564, tstat=-0.36, N=24 birds including 12 control birds).

Deaf and hearing birds exhibited different LO contingencies. While in hearing birds on average 46% of syllable renditions triggered light-off (p=0.08, tstat=-1.91, df=11, two-tailed t-test of hypothesis that LO rate is 50%), in deaf birds the average LO rate was 58% (p=0.04, tstat=2.45, df=9, two-tailed t-test of hypothesis that LO rate is 50%, Fig. 2g). Thus, only deaf birds diverged from the 50% expectation set by the previous day.

To analyze the sensitivity of birds’ vocal response irrespective of whether they were attracted or repelled by light-off, we quantified their magnitude pitch responses as the normalized pitch change per day (d’ value) aligned in the direction of global pitch change, implying that the average magnitude change was always a positive number. The daily magnitude pitch change was larger by 135% in deaf birds than in hearing birds (difference 0.16 d’/day, p=0.01, tstat=2.73, df=20, n=12 hearing and n=10 deaf birds, two-tailed t-test), Fig. 2f. Thus, visual feedback is much more salient when it substitutes auditory feedback in deaf animals than as a supplemental feedback in hearing birds.

Nevertheless, deaf and hearing birds similarly changed their songs in response to substitution, their pitch changes were confined to a temporal window within roughly 10 ms of the targeted time window for LO delivery, Fig. S2 and S3.

### The inversion in valence does not reflect a preference for darkness in deaf birds

A simple explanation of our findings could be that deafness induces an attraction to darkness for whatever reason. This explanation was ruled out after we replaced light-off by light-on stimuli and found strongly appetitive responses to such stimuli in deaf birds (0.71±0.07 d’/day in the direction of increasing light-on rate (0.61 and 0.77 d’/day), 2/2 birds significantly exceeded spontaneous pitch drift in control (unsubs) birds, p<0.05, two-tailed t-test on pitch changes per day, light-on contingency 77%±8% (75% and 79%), see Methods).

### A manipulation bonus seems to be required to explain valence reversal

Our vocal-light substitution paradigm forms a simple but powerful touchstone for theories of intrinsic motivation because 1) the vocal space of deaf birds is essentially binary (light on vs off), 2) the environment has no intrinsic dynamics (light only depends on pitch), 3) there is no evolutionary adaptation to LO stimuli, and 4) birds have no physiological need to sing a particular pitch (unlike their need of food intake). Despite this simple framework, most models of behavioral learning cannot accommodate valence inversion. In reinforcement learning (RL)^21^, stimuli have either appetitive or aversive effects and standard RL models cannot accommodate valence inversion for example via changes in baseline reward due to deafening^22,23^.

Our findings are also incongruent with computational models of directed exploration that involve an exploration bonus for action policies that are either informative^12–14^, diverse^8,9^, or simple^24^, Table 1. These theories have been designed to either improve the efficiency of reinforcement learning models or to model human behavior within a restricted class of multi-armed bandit problems^10,12,13^. In these models, agents choose actions that maximize the information gained about the environment, which is often modeled as an exploration bonus in proportion to the uncertainty of an action’s value^9,10^. Yet, in binary (and pitch-symmetric) worlds, knowledge gain is maximal when agents uniformly sample from their action repertoire, implying that such theories predict convergence of LO contingency to 50%^12,13,25^, which contradicts the divergence we found in subs birds.

**Table. 1:** Reinforcement learning characteristics and their compatibility with valence inversion.

| Reinforcement learning<br>model characteristic | Additional info | Can it explain<br>valence inversion? |
| --- | --- | --- |
| Changes in baseline reward | Caused by deafening | No |
| Exploration bonus | For informative actions | No |
|  | For diverse actions | No |
|  | For simple actions | No |
| Manipulation bonus | For impact | Yes |

In essence, to step above a LO contingency of 50%, a manipulation bias is required towards actions that impact the environment (such as light off). We introduced such a bias by defining a manipulation bonus *M*_*j*_ associated with action *j*. This bonus 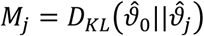 models the impact of action *j* in terms of the Kullback-Leibler divergence between the estimated sensory probability density 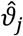 following action *j* and the same density 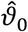 without any preceding action. Let us denote the LO probability following action *j* by 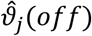 and the LO probability without acting by 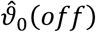. Because 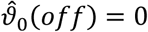, it follows that in deaf birds, the impact of action *j* is given by the Shannon surprise of light on: 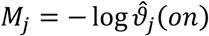. The impact is the larger the more likely the action triggers LO (the smaller 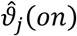). By experimental design, the impact is nonzero only for a small set of LO actions. An agent that maximizes impact will therefore exhibit a (manipulation) bias towards LO. In hearing birds, by contrast, the sensory environment includes vision and audition. Thanks to auditory feedback, all vocalizations in hearing birds elicit a nonzero impact. Thus, when hearing birds maximize impact, no particular action is singled out, which leads to absence of a manipulation bias towards light off.

In simulations, we modeled birds’ intrinsic reward *R*_*j*_ = *E*_*j*_ *+ M*_*j*_ *+ r*_*j*_ associated with action *j* as the sum of an exploration bonus *E*_*j*_ (given by information gain), a manipulation bonus *M*_*j*_, and an extrinsic punishment *r*_*j*_ *≤ 0* associated with light-off (*r*_*j*_ *< 0* only in case of light off), Fig. 3a, b. We simulated a simple agent that maximizes *R*_*j*_ using SARSA^26,27^, a standard RL framework. In simulations, we found that when the punishment term *r*_*j*_ per LO was such that deaf birds’ LO contingency converged to values above and hearing birds’ to values below 50%, Fig. 3d, the singing preference increased in deaf birds and it decreased in hearing birds, compared to their simulated controls, Fig. 3e, in excellent agreement with data. A manipulation bonus was required to reproduce these findings, Fig. 3d. Thus, when a behavioral goal is to detect impact via sensory feedback, such intrinsic reward can account for valence inversion and for high salience of substituted feedback. Furthermore, by design^28^, the model output in Fig. 3 agreed with known firing behavior of dopaminergic neurons^29^, which in hearing birds fire less than average on escape trials (no negative reinforcement) and more than average on hit trials (negative reinforcement), Fig. 3f.

**Fig. 3:**
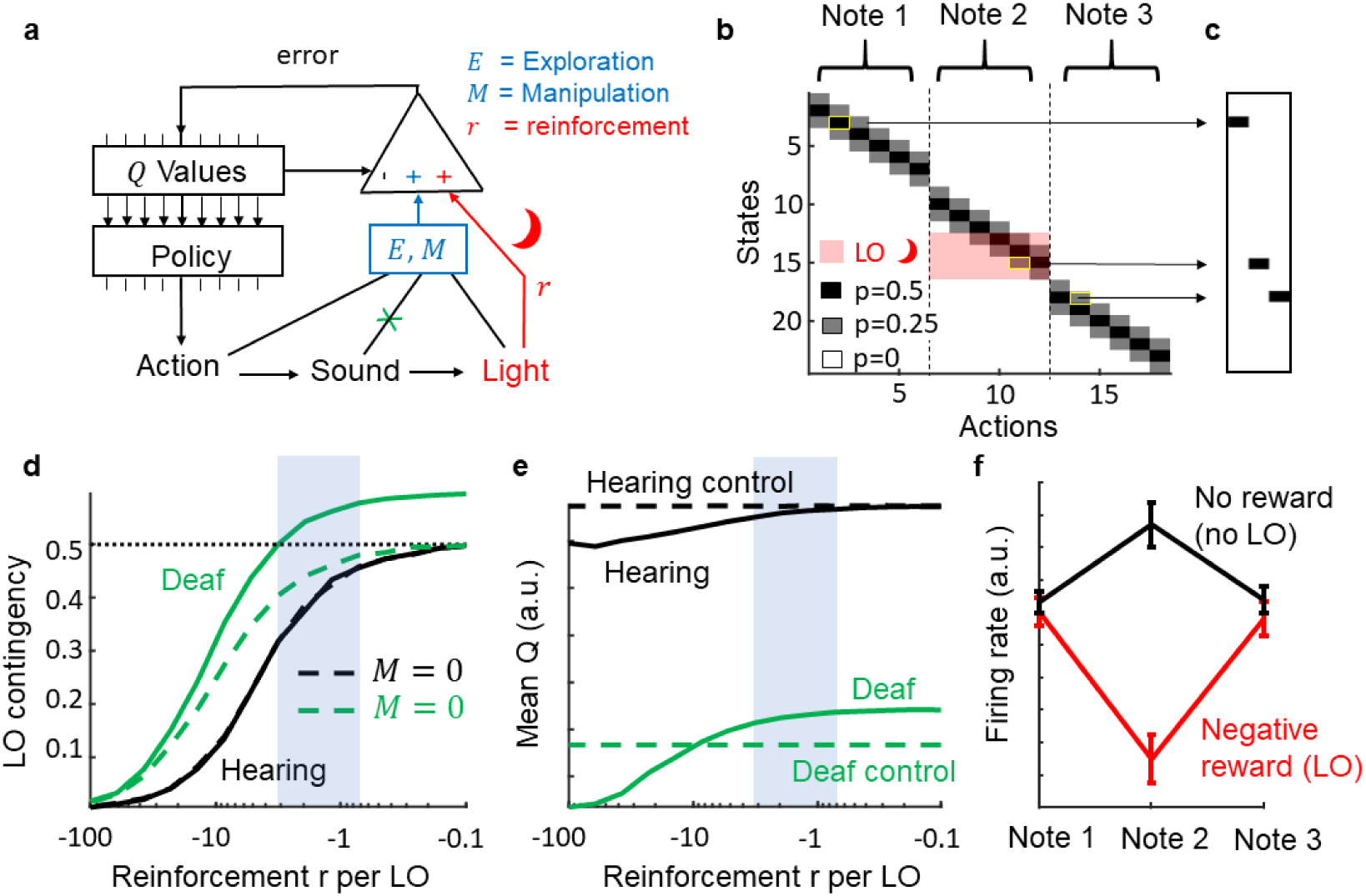
A manipulation bonus is compatible with valence inversion. **a**, We modeled a simple agent that maximizes total reward formed by the sum of the extrinsic reinforcement r (red), an exploration bonus E, and a manipulation bonus M given by impact. The agent’s greedy policy is to choose the action with maximal Q value (expected total reward). Deaf birds receive no auditory input (green cross). **b**, Markov model of an agent that generates one syllable composed of 3 consecutive notes, each associated with 6 possible actions. An action triggers one of three possible sensory states with probabilities depicted with gray shading. States 13-16 trigger light-off (red). **c**, Example syllable generated by the model (the underlying action-state pairs are delimited in yellow in **b**). **d**, Hearing birds trigger light-off on less than 50% of syllables for all choices of negative reinforcement r per LO. Deaf birds go above the 50% LO contingency (green). Without manipulation bonus (M = 0, dashed line) this is not the case (β = 0). **e**, Simulated subs birds are more motivated to sing than their controls (unsubs), the mean Q value (green, arbitrary units) for subs is above that of unsubs (dashed green). In hearing birds, the situation is reversed, they are less motivated than their controls. The blue dashed area indicates the plausible reinforcement-per-LO region that qualitatively matches our results. **f**, simulated firing rate of Area-X projecting dopaminergic neurons in hearing birds, modeled as reward prediction errors of a simple Q learning rule^26^. On aversively reinforced trials during Note 2 (modeling a brief LO or an acoustic white noise stimulus), the firing rate decreases (red), whereas on escape trials (no reinforcer, no LO), the firing rate increases (black). Error bars depict standard deviations (across simulated model birds).

### Basal ganglia lesions prevent visual reinforcement of pitch

Our reinforcement learning model suggests an involvement of the basal ganglia in mediating a manipulation bias. Dopaminergic neurons can drive selective pitch changes via their action in Area X^30–32^, a region homologous to the mammalian basal ganglia^18,33^. To probe for a manipulation bias in Area X, we made irreversible bilateral lesions in Area X of deaf birds. When these birds were subjected to substituted LO feedback, none of them (N=5) changed pitch in excess to deaf controls (p>0.05 for all birds, two-sampled t-test), Fig. 4a-c. One bird in which the lesion did not overlap with either Area X or LMAN in both hemispheres changed pitch significantly compared to deaf controls (p=0.01, two-sampled t-test). In lesioned subs birds, the magnitude of average pitch change per day was smaller than in unlesioned subs birds (difference −0.22 d’/day, p = 0.003, tstat=-3.62, df=12, two-tailed two-sample t-test), and the daily pitch change in lesioned subs birds was not significantly different from zero (−0.02 d’/day, p=0.70, SE=0.05, tstat=-0.39, df=97, n=5 birds, fixed effect), Fig. 4d. In combination, these findings show that Area X is necessary for expressing adaptive pitch responses to substituted feedback.

**Fig. 4:**
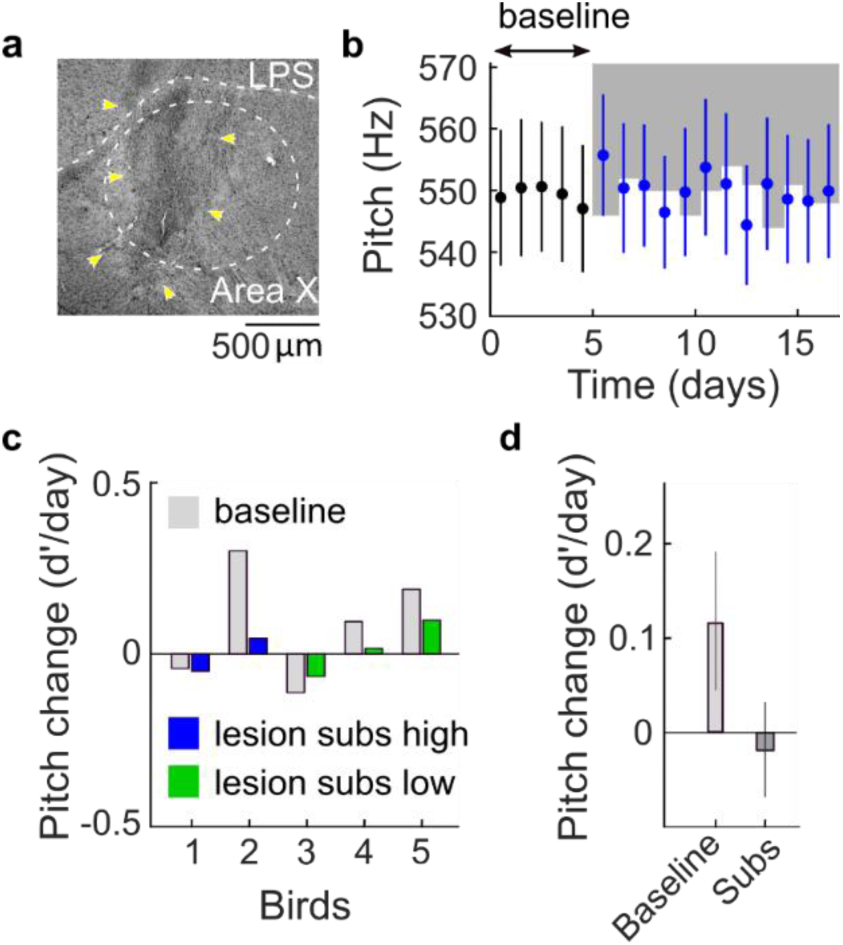
A basal ganglia pathway is necessary for adaptive responses to substituted feedback. **a**, Example sagittal brain section of a bird with lesion (yellow arrows) in Area X (dashed white ellipse). The lamina pallio-subpallialis (LPS) is indicated by the white dashed line. Anterior is towards the left, posterior to the right. **b**, Means (dots) and standard deviations (whiskers) of daily pitch distributions of an example deaf bird with bilateral lesions in Area X. There is no clear adaptive response to substitution. **c**, The average pitch changes (d’/day) in deaf birds with Area X lesions during baseline (light grey), during high-pitch substitution (blue), and during low-pitch substitution (green). **d**, The two fixed-effect terms of a mixed linear effect model and their standard errors: the daily pitch changes 1) during baseline (left bar, p=0.10), and 2) during substitution in the direction of increasing LO rate (right bar, p=0.70).

## Discussion

We found that elimination of auditory feedback induces appetitiveness of an otherwise aversive visual reinforcer of song. This finding helps to refine our understanding of the neural basis of vocal learning. Namely, because targeted changes of vocal skills can occur without hearing, it follows that evaluation of auditory performance is not a prerequisite for vocal plasticity in adulthood, unlike commonly assumed^29,33^. Our findings do not rule out the use of vocal performance for template-based song learning^34^, but they showcase that some forms of vocal learning do not rely on auditory representations of song, but rather on motor representations.

This means that the song system is able to assign a cross-modal signal such as light the role of an instructive signal that can selectively bias motor variability. To enable flexible assignment of visual signals (light brightness) to specific motor features (pitch), the visual system must feed into the song system in a computationally powerful way. We find that this cross-modal circuit involves the basal ganglia, which provides some clues as to the underlying neural mechanisms. For one, given that cerebral neurons efferent to the basal ganglia do not even respond to auditory feedback during singing^35–37^, it is unlikely that multimodal visual-vocal neurons are involved in this cross-modal learning flexibility. Rather, a large body of work on the basal ganglia suggests existence of an error-like signal that reinforces time-resolved motor representations of song^29,31,38^. Our work therefore suggests that the avian basal-ganglia part of the song system has evolved to support multimodal learning independent of the sensory modality of reinforcement.

We found that substitution uncovers a strong drive to manipulate. This drive better explains our observations than intrinsic motivations such as activity and exploration^15^. Normally, in healthy vocalizers, the need to manipulate is satisfied and does not constrain the brain’s valence system. However, when sensory feedback is lacking, this need becomes overwhelming to the point that it can override the valence even of aversive stimuli. This remarkable dictatorship of the manipulation drive emphasizes one of the most basic needs associated with motor actions, which is to perceive sensory feedback.

By design, the slightly aversive LO events are the only feedback that deaf birds experience, which is why to satisfy their manipulation drive, they prefer it over no response at all (‘something’ is preferable for a curious agent over ‘nothing’). One function of song in birds may thus be to exert an influence on the environment to signal the singer’s presence, even if confirmed only by visual feedback as in our experiment. Birds’ tendency to avoid overlap in their vocalizations^40^ is evidence for such a need of undisturbed signaling.

In humans, there exists a compelling analogy of the remarkable transformation of affective stimulus valence we observed. Namely, a manipulation drive shows up during boredom, which can prompt subjects to display behaviors that evidence paradoxical preference of otherwise aversive stimuli^41,42^. Lacking an alternative, subjects prefer an unpleasant experience rather than none at all.

Regarding substitution as a therapy, perhaps the frustration experienced by users of substitution devices is linked with the level of uncontrollability of the substituting input. Our findings suggest that users of substitution devices might avoid motor actions unless these elicit some form of substituting sensory feedback. Although the intention behind a substitution system usually is to maximize the conveyed information, to merely maximize information can conflict with a manipulation bias that may draw the use of a device away from its intended purpose.

We suspect that substitution devices are preferred when they provide feedback about motor actions, making subjects feel empowered through the new sense. By designing substitution systems as part of closed sensorimotor loops^4^, the systems may stimulate motor learning, which can be fun as in tennis practice or piano playing rather than strenuous as in learning a foreign language (analogies between learning to make use of substituted input and reading have been drawn^1^). Perhaps, acceptance of substitution devices would also increase when training setups are designed to let subjects predictably manipulate the substituting sensory input, in line with insights from interviews conducted with users of substitution devices^43^.

To focus on manipulation biases in substitution therapies must not necessarily interfere with information gain; rather, such biases may be exploitable on their own when they point towards desirable actions. Binary feedback signals are known to promote robust motor learning^44^. For example, the blind might benefit from a signal that reports the inverse distance of a hand to an object of choice. Or, in the context of speech rehabilitation, the hearing impaired may benefit from short-latency feedback when their variable speech agrees with signals of high comprehensibility; such feedback could be provided as visual signal (e.g. displayed in augmented reality devices) or as vibro-tactile signal^45^.

Manipulation biases might also be relevant in neuroprosthetic systems that aim to increase the perceptual space of subjects. In sensory neuroprosthetics, the sensor is not substituted but bypassed by electrical stimulation of downstream neurons. While neuroprosthetic closed-loop systems have only recently started to be explored in the sensory domain^46^, closed-loop systems are very common in the motor domain^47^, where animal models have played a crucial role in the development of a wide range of those systems^47,48^. Closed-loop motor systems achieve better performance than open-loop systems^47,48^ and there is a distinct performance benefit of high feedback rates^49^. These facts lend support to the idea that sensory neuroprosthetic systems will also benefit from closed-loop design. The zebra finch as an animal model may lend itself to exploring the closed-loop benefits of sensory neuroprosthetics.

Key to our findings is that songbirds have a strong manipulation drive. Abstracted as a principle of software agents, such a drive can serve some computational functions. Conceptually, manipulation seeking can be preferable to knowledge seeking because the latter is uninformative about relevance. For example, a manipulation drive can prevent an agent from getting stuck in front of a computer screen displaying random stimuli, which would otherwise be the most absorbing stimulus for a purely knowledge-driven agent that does not distinguish between self-generated and external stimuli. It is therefore not surprising that concepts such as manipulation and impact are gaining in importance in machine learning. In a recent curiosity-driven RL approach, it was found that a focus of actions on self-generated sensory feedback can dramatically expedite learning progress^50^.

Further impulses for understanding the motivational drives of spontaneous behaviors are highly needed. We propose that sensory substitution is a promising paradigm not just to experimentally characterize the motivation to manipulate, but also to dissect the neural representations of affective valence^51^ and to probe how substituting input is integrated into an existing circuit on the level of single cells, which so far is only understood on the level of brain areas^1,39^. Because the manipulation drive seems to have access to cross-modal learning mechanisms that are as fast and efficient as those of normal motor learning, sensory substitution and the manipulation drive it reveals may provide a glimpse on some of the enabling factors of evolutionary adaptations.

## Acknowledgments

We thank Florian Engert, Maneesh Sahani, and Georg Martius for helpful discussions and Benjamin Grewe, Thomas Tomka, and Catherine del Negro for helpful comments on the manuscript. Funding support was provided by the Swiss National Science Foundation (Project 31003A_182638). The authors declare no competing financial interests.

Data and scripts can be accessed at the ETH Research Collection.

## Author Contributions

Conceptualization, A.T.Z. and R.H.R.H.; Formal Analysis, A.T.Z. and R.H.R.H.; Investigation, A.T.Z. and S.C.L., Data Curation, A.T.Z. and M.R.; Writing – Original Draft, A.T.Z. and R.H.R.H.; Writing – Review & Editing, A.T.Z., N.G. and R.H.R.H.; Supervision, N.G. and R.H.R.H.

## Materials and Methods

### Subjects and song recordings

We used 50 adult male zebra finches (*Taeniopygia guttata*) raised in our breeding facilities in Zürich (Switzerland) and Orsay (France). At the beginning of the experiment, birds were between 90 and 200 days old. During the experiment, birds were housed individually in sound-attenuating recording chambers on a 14/10 h day/night cycle. Access to food and water was provided *ad libitum*. After 2-5 days of familiarization in the experimental environment, birds resumed singing at a normal rate. Songs were recorded with a wall-attached microphone, band-pass filtered, and digitized at a sampling rate of 32 kHz. All experimental procedures were approved by the Veterinary Office of the Canton of Zurich and by the French Ministry of Research and the ethical committee Paris-Sud and Centre (CEEA N°59, project 2017-12).

### Visual substitution of pitch

To provide pitch substitution, we ran a custom LabVIEW (National Instruments, Inc.) program. We targeted a harmonic syllable using a two-layer neural network trained on a subset of manually clustered vocalizations. We evaluated pitch (fundamental frequency) in a 16-ms window at a fixed delay after the syllable detection point (which occurred at a roughly constant time lag after syllable onset). We estimated pitch using the Harmonic Product Spectrum algorithm^52^, our code is published at the ETH Research Collection, also in the Matlab (Mathworks Inc.) language.

We provided pitch substitution by switching off the light (using a relay) in the sound recording chamber after a delay of 12 ms and for a duration in the range of 100 to 500 ms. Two birds were put in dim light and substitution was provided turning on an additional light instead of switching off the light. We substituted either high or low pitch, depending on a manually set threshold.

To cumulatively drive pitch of the targeted syllable away from baseline, every morning, we adjusted the pitch threshold to the median pitch value from the previous day based on all noncurated neural network detections (in 24/328 days from 15/29 birds, we did not set the threshold to the median value because of a software crash on the previous day). In 15 birds (6 hearing and 9 deaf, among which2 were Area X lesioned birds and 1 missed lesion) we delivered substitution on high-pitched syllable renditions, and in 15 birds (6 hearing and 9 deaf, among which 3 were Area-X lesioned birds) on low-pitched syllable renditions. **Subs birds** were deaf birds with LO substitution, **unsubs birds** were deaf and unsubstituted birds; **LO birds** were hearing and subjected to LO, and **no LO birds** were hearing and not subjected to LO.

### Surgeries

Before the onset of surgery, we provided analgesia with the nonsteroidal anti-inflammatory drug carprofen (2-4 mg/kg, Norocarp, ufamed AG, Sursee, Switzerland) given intra muscularly (IM). Birds were deeply anesthetized using isoflurane (1.5-3 %) and placed in a stereotaxic apparatus. We applied the antiseptic povidone-iodine (Betadine, Mundipharma Medical Company, Basel, Switzerland) to the skin at the incision site, followed by the local anaesthetic lidocaine in gel form (5 %, EMLA, AstraZeneca AG, Zug, Switzerland).

### Deafening procedure

In the stereotax, the head angle formed by the flat part of the skull above the beak and the table was set at 90°. The skin was opened above the hyoid bone and the neck muscles were gently pushed back to expose the semi-circular canals. A hole was made in the skull to access the inner ear below the semi-circular canals. The cochlea was visually identified based on the surrounding bone structure and a small hole was made with forceps into the cochlear base. We removed the cochlea from the cavity with a custom-made tungsten hook and took a picture of both intact cochleae including the lagenas to document the success of the procedure, Fig. 1b.

### Area X lesions

We set the head angle formed by the flat part of the skull above the beak and the table to 35° and drilled a window into the skull above Area X. Area X was localized based on stereotaxic coordinates and identified through presence of tonically firing neurons, recorded with a 0.6-1.7 MΩ tungsten electrode attached to a vertical manipulator. We injected in each hemisphere 1 μl of ibotenic acid (Tocris) near the center of Area X. Injection sites were located on average 1.5-1.9 mm medial-lateral (ML), 5.5-6.0 mm anterior-posterior (AP), and 2.8-3.5 mm dorsal-ventral (DV) from the bifurcation of the midsagittal sinus (lambda). Injections were performed using a borosilicate glass pipette (BF-120-69-10, Sutter instrument) pulled with a Picospritzer (Parker Inc.) and broken with forceps to a tip diameter of about 10 μm.

### Histology

At the end of the experiment, birds were euthanized with an overdose of intraperitoneal injection of sodium pentobarbital (200 mg/kg, Esconarkon, Streuli Pharma AG, Uznach, Switzerland) and intracardially perfused with 4% paraformaldehyde (PFA) before brains were removed for histological examination. Brains were rinsed in a 0.01 M phosphate buffer solution. The hemispheres were separated from each other, glued on a metal plate, and embedded in 3% agar. Sagittal slices of 80 μm thickness were cut with a Thermo Microm HM650V microtome and mounted on slides for Nissl staining.

### Statistical pitch analysis

We curated the neural network detections manually by visually removing misdetections (detection of noises or timepoints in the song that did not correspond to the selected target syllable). We quantified the effects of light-off (LO) on pitch using **d-prime values** 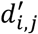:

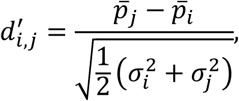

where 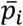 and 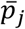 are the mean pitches on days *i* and *j*, and 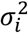 and 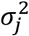 are the variances, respectively.

### LO start criterion

LO started after at least 5 days of stable singing (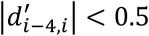 with *i* being the last day of baseline); in two birds, LO started earlier and in one bird, LO started later because of technical issues and unforeseen scheduling constraints. In deaf (subs) birds, LO started on average 15 days after deafening (range 7 to 33 days) and in hearing (LO) birds it started after at least 7 days in isolation.

### LO end criterion

We ended the LO paradigm when the absolute mean pitch change (relative to baseline) either exceeded at least 2.5*d’* or when it stabilized near zero, which was defined as 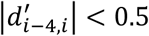 with the index *i* referring to the last day of light off (in one bird, we ended substitution before this criterion was reached because the song degraded too much for reliable syllable detection). The duration of the substitution paradigm did not differ significantly between birds in the hearing and deaf groups (p = 0.39, two-tailed two-sample t-test, mean hearing = 13 days, mean deaf = 11 days). Thus, the observed differences between hearing and deaf birds in Fig. 2e, f were not due to differences in time spent in the experimental chamber.

The **average daily pitch change** 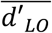 **during *substi*tution** in each animal (Figs. 1g and 2c) we quantified as 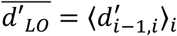, where the angle brackets denote averaging across all days *i* with LO (starting from the second day).

Similarly, the **average daily pitch change** 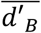 **during the baseline period** in each animal (light grey bars in Figs. 1g and 2c) we quantified as 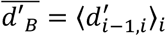, where the average runs across the last 4 days *i* before LO.

### Magnitude pitch change

We assessed the magnitude pitch change in each bird irrespective of its preference (attraction or repulsion by LO). To discount for preference, we first defined the **global direction** *δ* **of pitch change** during substitution as 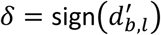, where *b* is the last day of baseline and *l* is the last day of LO exposure (*δ* corresponds to the direction of the colored bars in Figs. 1g and 2c). Thus, if birds shifted pitch upward towards higher values, *δ* = *1*, and if birds shifted pitch down, *δ* = −*1*. In each animal, we computed the **mean aligned pitch change** 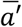 during substitution as the average daily change 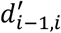 multiplied with 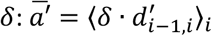, (*i*=6, …, end). Fig. 2f shows 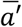 averaged over all birds. For control birds (unsubs, no LO), the direction of change *δ* was calculated analogously.

### Sound features other than pitch

To test whether substitution-induced sound feature changes of the target syllable were specific to pitch, we inspected sound features including syllable duration, amplitude modulation (AM), frequency modulation (FM), and entropy. Syllable duration was defined by the interval between consecutive threshold crossings of RMS sound waveform, the threshold for each animal was kept constant for all days analyzed. AM, FM, and entropy were computed as means over the entire syllable. For the analysis, we combined low-pitch and high-pitch substituted birds, all feature values in low-pitch birds were first multiplied by −1 to account for the anti-symmetry between their treatments. As a group, we compared the feature *d′* values between the last LO day and the last day of baseline (paired two-tailed t test), Supplementary Fig. S1a.

### Control birds

To evaluate whether an individual bird responded to substitution, we compared the daily pitch changes 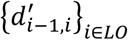 of its targeted syllable to daily pitch changes of harmonic syllables (containing a harmonic part longer than 100 ms) in many control birds (not exposed to LO). In subs birds, the control group was formed by 12 harmonic syllables in 9 deaf animals, and in LO birds, the control group was formed by 12 harmonic syllables in 12 hearing birds. To account for possible pitch drifts caused by deafening or by time spent in the experimental chamber, the time window for pitch analysis in unsubs birds was matched to the substitution period in the subs bird, i.e., the first day analyzed in control birds occurred at the same time lag after deafening as the first LO day. Also, the number of days analyzed was identical in the subs bird and in control birds (same for LO and no-LO birds). We paired a subs bird only to controls that produced more than 100 song motifs on each day during the matched time periods. Two unsubs birds had to be excluded because they produced fewer than 100 renditions on days 11 and 12 after deafening (these birds could not be time-matched to any subs bird, resulting in 10 harmonic syllables from 7 control birds).

### Pitch response testing

To test whether an individual bird significantly changed its pitch in response to LO, we compared all its daily pitch changes during LO to all daily pitch changes in control birds in matched time windows (at significance level p=0.05, two-tailed two sample t-test, indicated by asterisks in Figs. 1g, 2c).

For the population analysis, we compared daily pitch changes in all subs/LO birds against all unsubs/no-LO controls. We randomly paired 10 syllables in control birds (dark grey bars in Fig. 1g) with the 10 syllables in subs birds (under the constraint that analysis days could be temporally matched). The pairing is depicted in Fig. 1g such that target syllable 11 was paired with subs bird 1, target syllable 12 with subs bird 2, etc. We did the same for the 12 syllables in LO birds in Fig. 2c, i.e., target syllable 13 was paired with LO bird 1, target syllable 14 was paired with LO bird 2, etc. All pairings were time-matched, i.e., the early (baseline, light gray bars in Fig. 1g) and late time periods in controls were defined according to the baseline and LO periods in the treated bird.

### Linear mixed effect model

To test whether subs/LO birds exhibited a common direction of pitch change (either towards LO or away from it), we modeled daily pitch changes 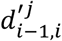 in bird *j* in response to LO with a linear mixed effect model:

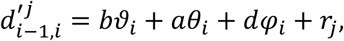

where the three fixed effect terms *b*, a, and *d* common to all birds were: the daily pitch change *b* during baseline (*ϑ*_*i*_ = *1* if day *i* is during baseline and *ϑ*_*i*_ = *0* otherwise), the pitch drift a without LO (*θ*_*i*_ = *1* in control birds if days *i* and *i* − *1* occurred after baseline and *θ*_*i*_ = *0* otherwise), and the daily pitch change *d* caused by LO (*φ*_*i*_ = *1* for LO high and *φ*_*i*_ = −*1* for LO low birds, provided both days *i* − *1* and *i* were LO days). The *r*_*j*_ are zero-mean Gaussian noise terms that account for variability among birds. We separately fitted linear mixed effect models to deaf and to hearing birds.

When we reduced the model to two fixed effects (combining the terms *b* and *a* into a single term describing spontaneous pitch drift during both baseline in LO-treated birds and all days in control animals) we found the results displayed in Fig. 1h and Fig. 2d to be qualitatively unchanged.

### Song degradation

To assess song degradation caused by deafening (Supplementary figure S1b-e), we inspected non-targeted syllables, comparing renditions at the beginning and the end of the experiment. Tschida and Mooney showed that both entropy and entropy variance significantly change after deafening^20^. Mean entropy is a measure of syllable noisiness and variance entropy of syllable complexity. To follow suit and inspect mean and variance entropy, we first semi-automatically clustered all (non-targeted) syllables using a nearest neighbor approach in the spectrogram domain. We only considered syllables that were sung more than 100 times on each day (21 syllables in hearing birds and 18 syllables in deaf birds). We calculated for each syllable type the magnitude mean-entropy change as the absolute difference in mean entropy between the last day before deafening and the first day after LO ended. For hearing birds, we chose the first day analyzed such that the duration of the analysis window matched the window in deaf birds. As a result, the intervals between the first and last day of the experiment did not significantly differ between birds in the hearing and deaf groups (p = 0.86, two-tailed two-sample t-test, mean hearing = 32 days, mean deaf = 31 days). Thus, differences between hearing and deaf birds in Supplementary Fig. S1b-e were not due to differences in time spent in the recording chamber.

In agreement with Tschida and Mooney, we found that deaf birds have a larger magnitude variance-entropy change than hearing birds (difference 0.32, p = 9.3*10^−4^, tstat = 3.60, df = 37, two-tailed two-sample t-test, Suppl Fig. S1c). However, we found no difference in magnitude mean-entropy change (p=0.61, Suppl Fig. S1b). Note that Tschida and Mooney did not perform time-matched comparisons against a group of hearing birds as we did, but they compared entropy to baseline measurements taken before deafening, implying that mean entropy changes in their study could have been caused by birds’ gradual adaptation to the recording chamber, irrespective of the deafening procedure.

For non-targeted syllables, we calculated the pitch coefficient of variation *CV*_*i*_ on day *i* as 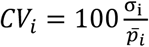 As we had done for targeted syllables, we calculated the pitch within a fixed 16 ms window during a harmonic part of the syllable (provided the latter existed, i.e., a harmonic part was found in 9/18 syllables in deaf animals and in 9/21 syllables in hearing animals). The difference between the coefficients of variation on the last day of deafening and on the first day after LO is shown in Suppl Fig. S1d. In deaf birds, this difference was larger than in hearing birds.

To compute spectral changes due to deafening we performed a bias-variance decomposition. To calculate spectrograms, we first tapered the sound waveform using a Hamming window of 512 samples. The windowed signal was transformed into a linear-power sound spectrogram using the discrete fast Fourier transform computed over segments of 512 samples and nonoverlaps of 128 samples (corresponding to 4 ms). The log-power sound spectrogram was then obtained by taking the natural logarithm of the linear-power sound spectrogram after adding an offset of 0.1 (corresponding roughly to the 75^th^ percentile). We computed the spectrograms of non-targeted syllables within a time window defined by the duration of the shortest syllable rendition. To achieve robustness to low-frequency noise present in the recordings, we ignored the lowest 10 frequency bins corresponding to a frequency cutoff at 625 Hz. The spectrogram bias of a particular syllable was defined as the Euclidean distance between the average spectrograms on two separate days: on the last day before deafening and the first day after the end of the LO period. The spectrogram variance was defined as the average pixel-wise variance on a given day. There was no significant difference between hearing and deaf birds in terms of either spectrogram bias or variance (bias: p=0.45, tstat=0.77, df=39, variance: p=0.32, tstat=-1.01, df=39, two-tailed two-sample t-test, Suppl Fig. S1e and f). Thus, the substitution period was too short to lead to a major spectral song degradation.

### Localization of pitch changes

Next we assessed the temporal dynamics of pitch changes in response to light off. We computed pitch traces over the entire syllable in a sliding window of 16 ms and plotted their temporal statistics at a time resolution of 1 ms, supplementary Fig. S2. In each bird, to compare pitch traces from the last day of light off with traces from the last day before light off, we computed d’ values between the two distributions at 1-ms time scale relative to the window of LO delivery, Supplementary Fig. S3. For bird 1, the syllable detection point was not stable and over the experiment the pitch window slightly shifted relative to syllable onset (see Supplementary Fig S2a, bird 1). To avoid the visualization of spurious pitch changes due to syllable detection jitter, we corrected for this temporal jitter before plotting pitch difference traces across the LO period in individual birds and their averages, Supplementary Fig. S3 (we aligned all syllables to their onsets and defined the pitch window at the mean time lag of all windows on the day before light off).

