## Supplementary Material for "Sensory substitution reveals a manipulation bias"

**Reinforcement Valence in Markov Decision (MD) Models**

We describe a computational model of spontaneous behavior wherein a singing bird behaves like a simple probabilistic agent. We express the model in a reinforcement learning (RL) framework, according to which birds try to maximize the total reward obtained from their songs. This reward encompasses both extrinsic and intrinsic components, the latter of which we try to estimate.

### 1. Markov Modeling Framework

We assume that singing forms a Markov process in which at time $t$ $(t = 1, \dots, T)$, the **motor action** $a^t$ triggers the **sensory state** $s^t$ according to the unknown **Markov observation matrix** $\vartheta_j(k) = P(s^t = k | a^t = j)$ which denotes the probability of observing sensory state $k$ given motor action $j$. One can think of this observation matrix as modeling the vocal organ including the motor noise associated with singing. In the following, $k$ is the index of all distinct sensory (visual and auditory) states, and $j$ is the index of all distinct motor states.

Through song practice, the bird obtains information about this observation matrix. We assume it does so by keeping track of how often it chose a given action and how often it observed a given state as a result. When at time $t$, the bird produces action $j^*$ and observes state $k^*$, the bird's **action-state counter** $C_{j,k}^{t}$ is updated as follows:

$$C_{j,k}^{t} = \begin{cases} \tau \cdot C_{j,k}^{t-1} + 1 & \text{for } k = k^* \text{ and } j = j^* \\ C_{j,k}^{t-1} & \text{otherwise} \end{cases}$$

Here, $\tau$ models a forgetting rate to mimic birds' limited counting ability. We assume that forgetting is triggered by actions, not by time, to agree with one-shot aversive conditioning that can be long lasting[1]. Based on this action-state counter $C_{j,k}^{t}$, we want to formulate an intrinsic reward such that a greedy bird will choose actions in agreement with the behavior of both hearing and deaf birds in our experiments.

### 2. Song Model

For simplicity, we assume the bird sings a motif composed of a single syllable which in turn is composed of three **notes** $i = 1,2,3$. Because adult zebra finch syllables are very stereotyped, these notes are always produced in the same order. The part of motor variability that is voluntarily controlled by the bird arises from each note having a distinct set of six actions associated with it. These actions span diverse note variants that are distinct in terms of their sound features including pitch. Thus, the bird's motor repertoire is composed in total of 18 different **actions** $a_j$ $(j = 1, \dots, 18)$, where actions 1-6 are associated with note $1$, actions $7 - 12$ with note $2$, and actions $13 - 18$ with note $3$.

To model motor noise that the bird cannot voluntarily control, we assume that each action probabilistically leads to one of three different sensory states, as depicted in Fig. 3B. The sensory states represent the actual sounds being produced. The center state has high probability (0.5) and the two flanking states have lower probabilities ($0.25$ each). The sensory states associated with a note tile the possible pitch values in a topographical manner such that a given state can be elicited by up to three different actions (because of motor noise). Without loss of generality, we assume that the sensory spaces of all three notes are distinct and nonoverlapping (complex syllable), which means we are dealing with a total of $25$ different sensory states $s_k$ $(k = 1, \ldots, 25)$, where states $1-8$ are associated with note $1$, states $9-16$ with note $2$, states $17-24$ with note $3$ and the $25^{\text{th}}$ state corresponds to the silent state which hearing birds can never reach during singing.

In summary, when the bird sings note $i$, the **Markov observation matrix** $\vartheta_j(k)$ is given by

$$\vartheta_j(k) = \begin{cases} 0.5 & \text{for } k = k_0 \\ 0.25 & \text{for } k = k_0 - 1 \text{ or } k = k_0 + 1 \\ 0 & \text{otherwise,} \end{cases} \qquad (1)$$

where $k_0 = j + 1 + 8(i-1)$ is the index of the center state associated with the $j$th action. The roughly diagonal structure of the Markov observation matrix (Fig. 3B) amounts to assuming that the sensory and motor neural representations are well aligned with one another, which is the case for example when the brain forms a forward or inverse model of the vocal apparatus[2].

In note 2, half of the states (states $13-16$) trigger a brief LO event. The difference between hearing and deaf birds is that hearing birds experience all $24$ sensory states (corresponding to different pitch values and light conditions) whereas deaf birds experience only two states (light off and neutral).

### 3. Intrinsic Reward from Impact

We write the **total reward** $R_{j^*}^t$ at time $t$ and associated with action $a^t = j^*$ and observed state $s^t = k^*$ as the weighted sum of three terms: an exploration bonus $E_{j^*}^t$, a manipulation bonus $M_{j^*}^t$, and an extrinsic punishment $r^t$ associated with the lighting condition:

$$R_{j^*}^t = E_{j^*}^t + M_{j^*}^t + r^t. \tag{2}$$

The three main terms are:

A. The **exploration bonus** $E_{j^*}^t$ is defined as the Markov information gain defined as the log Kullback-Leibler (KL) divergence $D_{KL}\left(\hat{\vartheta}_{j^*}^{t-1} || \hat{\vartheta}_{j^*}^t\right)$ between $\hat{\vartheta}_j^t(k)$ and $\hat{\vartheta}_j^{t-1}(k)$[3,4]:

$$E_{j^*}^t = \log D_{KL}\left(\hat{\vartheta}_{j^*}^{t-1} || \hat{\vartheta}_{j^*}^t\right), \text{ with } \; D_{KL}\left(\hat{\vartheta}_{j^*}^{t-1} || \hat{\vartheta}_{j^*}^t\right) = \sum_k \hat{\vartheta}_{j^*}^{t-1}(k) \log\left(\frac{\hat{\vartheta}_{j^*}^{t-1}(k)}{\hat{\vartheta}_{j^*}^t(k)}\right).$$

The agent's (biased) **estimate** $\hat{\vartheta}_j^t(k)$ of the Markov observation matrix $\vartheta_j^t(k)$ is given by the fraction of times a given state has been observed, $\hat{\vartheta}_j^t(k) = \frac{1+C_{j,k}^t}{1+\sum_l C_{j,l}^t}$. The exploration bonus is large when the bird explores rare actions, in which case the difference between the action-state counters $C_{j,k}^t$ and $C_{j,k}^{t-1}$ is large and so is the distance $D_{KL}\left(\hat{\vartheta}_{j^*}^{t-1} || \hat{\vartheta}_{j^*}^t\right)$ between the estimated observation matrices $\hat{\vartheta}_{j^*}^{t-1}$ and $\hat{\vartheta}_{j^*}^t$.

B. The **manipulation bonus** $M_{j^*}^t$ associated with action $j^*$ is defined as:

$$M_{j^*}^t = D_{KL}\left(\hat{\vartheta}_0^t || \hat{\vartheta}_{j^*}^t\right),$$

where $D_{KL}$ denotes the KL divergence $D_{KL}\left(\hat{\vartheta}_0^t || \hat{\vartheta}_{j^*}^t\right) = \sum_k \hat{\vartheta}_0^t(\mathrm{k}) \log\left(\frac{\hat{\vartheta}_0^t(k)}{\hat{\vartheta}_{j^*}^t(k)}\right)$. In the context of our experiments, the light-off state can only be triggered by singing and thus $\hat{\vartheta}_0(k = \text{light off}) = 0$. When we set $\hat{\vartheta}_0^t(k = l\text{ight off}) = 0$ in the definition of the manipulation bonus, we obtain $M_{j^*}^t = -\log\left(\hat{\vartheta}_{j^*}^t(k = \text{light on})\right)$, i.e., the deaf bird will try to maximize the surprise of the light-on state by triggering LO as often as possible.

C. The external reward $r^t$ is zero most of the time and has a fixed negative value $\mathscr{r} < 0$ when the light goes off (because bird's vision is obstructed):

$$r^t = \begin{cases} \mathscr{r} & \text{if note } i = 2 \text{ and } k^* \in \{13, \dots, 16\} \\ 0 & \text{otherwise.} \end{cases}$$

External rewards are maximized when the bird avoids actions associated with light off.

**4. Action Choices in Reinforcement Learning Framework**

We express the model in a simple SARSA framework [5] in which the agent makes greedy action choices

$$j^* = \underset{j \in \text{ note } i}{\operatorname{argmax}} Q_j^t, \tag{4}$$

by maximizing an **action-value function** $Q_j^t$ that is defined as the running average reward [6]:

$$Q_{j^*}^t = Q_{j^*}^{t-1} + \alpha\left(R_{j^*}^t - Q_{j^*}^{t-1}\right), \tag{5}$$

with $\alpha$ being the learning rate. Initially, the action-value function $Q_j^t = 0$ is set to zero for all states $j$.

**5. Intuition**

In our stationary world, $\hat{\vartheta}_0^t(k = \text{light neutral}) = 1$ and the impact is maximized for actions $j^*$ for which $\hat{\vartheta}_{j^*}^t(k = \text{light off}) = 1$, i.e., when the bird turns the light off. Maximizing the total expected reward $R_{j^*}^t$ leads to a less radical action choice, thanks to the exploration bonus. In practice, birds find some balance between impacting the world and probing diverse actions.

**6. Model Details**

To produce Fig. 3 and Fig. 4, we used the following simulation parameters: $T = 600$, $\tau = 0.99$, and $\alpha = 0.05$. In Fig. 3C, we plotted the fraction of syllables that triggered LO as a function of the negative extrinsic reinforcement $r$ per LO. In Fig. 3D, we plotted the average action value $\langle Q_j^t \rangle_{T,j} = \frac{1}{18T} \sum_{j,t} Q_j^t$ for hearing and for deaf birds in the conditions with and without LO. We interpret high average action values as highly motivated birds that sing frequently and low action values as lowly motivated birds that sing infrequently.

For the simulation of dopaminergic neurons in Fig. 4, we defined the firing rate $f$ to be proportional to reward prediction error[7]: $f \propto R_{j^*}^t - Q_{j^*}^{t-1}$.

**Remarks**

- Our definition of the manipulation bonus $M_{j^*}^t$ that we termed impact is inspired from[8], where impact is used as a norm for characterizing people's question-asking strategies.
- Except for motor noise inherent in the definition of $\vartheta_j(k)$, all aspects of the agent are deterministic.
- In simulations, $R_{j^*}^t$ should be non-positive, otherwise simulated birds get stuck on the one action for which the total reward is most positive. The logarithm in the definition of the exploration bonus tends to make sure that $R_{j^*}^t$ remains negative.
- None of the simulation results critically depends on the number of actions (i.e., six), the number of sensory states (i.e., eight) per note, and the amount of motor noise modeled. By contrast, it is important that the Markov observation matrix $\vartheta_j(k)$ in Equation (1) significantly differs between

hearing and deaf birds (we assumed it is roughly diagonal and 24x18-dimensional in hearing birds in contrast to degenerate and 2x18-dimensional in deaf birds).

- The exploration bonus $E_{j^*}^t$ is not strictly needed in simulations. In a much simpler version of our model, the same qualitative results apply when $E_{j^*}^t$ is set to a fixed value, e.g. -6. The fixed subtraction of -6 on every chosen action in combination with Equation (4) guarantee that the bird will probe diverse actions, akin to maximizing the exploration bonus.
- Fig. 4 should be interpreted as a model of dopaminergic neuron firing reported in[9]. In principle, the only necessary term in Equation (4) to model results in[9] is $r^t$, which corresponds to the negative reward associated with white noise sound bursts (instead of LO). In this respect, the bonuses $E_{j^*}^t$ and $M_{j^*}^t$ mainly contribute variance.

Although our model is expressed as a model of dopaminergic neurons, it is not meant to imply that all 3 terms in Equation (4) must necessarily be signaled by dopaminergic neurons. It is conceivable that other neuromodulators such as serotonin have the function of signaling intrinsic rewards.

### Supplementary Figures: S1-S3

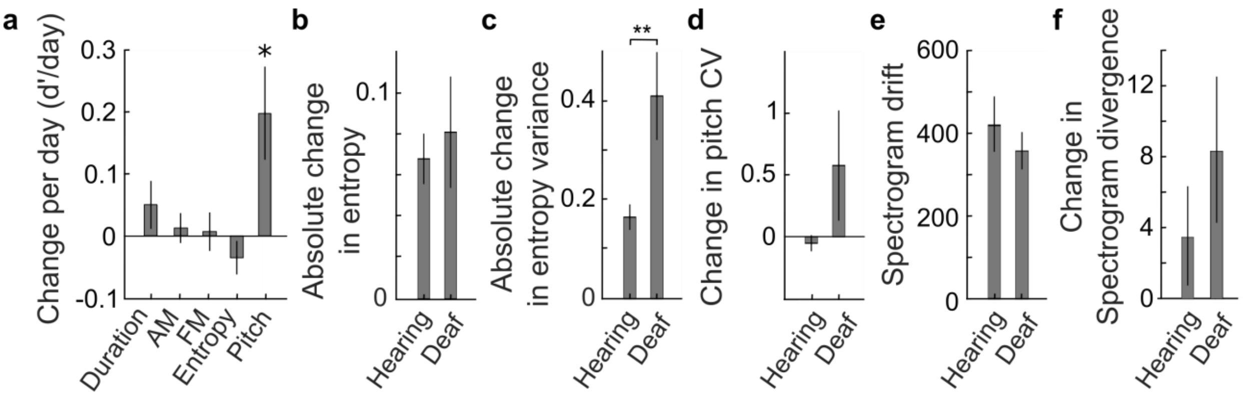


***Fig. S1: Syllable dynamics in hearing and in deaf birds. a,*** *In deaf birds, pitch-contingent LO induced changes in pitch, but no other sound feature of the target syllable. Shown are changes (d' per day) between the first and the last day of brief LO exposure for the following features: syllable duration, amplitude modulation (AM), frequency modulation (FM), entropy, and pitch. Bars indicate averages across birds and whiskers the standard errors of the mean. Before averaging, feature changes in LO low birds were multiplied by a factor -1 to align the changes with those of LO high birds. Significant nonzero changes were seen in pitch (p=0.044, two-tailed t-test, df=9, tstat=0.45, sd=1.14), but no other feature (p>0.05, two-tailed t-test). To compute d' per day in each animal, we divided the changes between the first and last day by the number of intermittent days.* ***b-e,*** *Non-targeted syllables diverged more in deaf birds than in hearing birds. In deaf birds, the bars indicate the mean syllable changes from the last day before deafening to the first day after LO, in hearing birds, changes are measured across equal time intervals (on average 32 days). Shown are* ***b*** *entropy,* ***c*** *entropy variance,* ***d*** *pitch coefficient of variation (CV),* ***e*** *drift in mean spectrogram,* ***f*** *spectrogram divergence. The error bars indicate the standard errors of the means and the stars indicate p value ('**': p=0.01).*

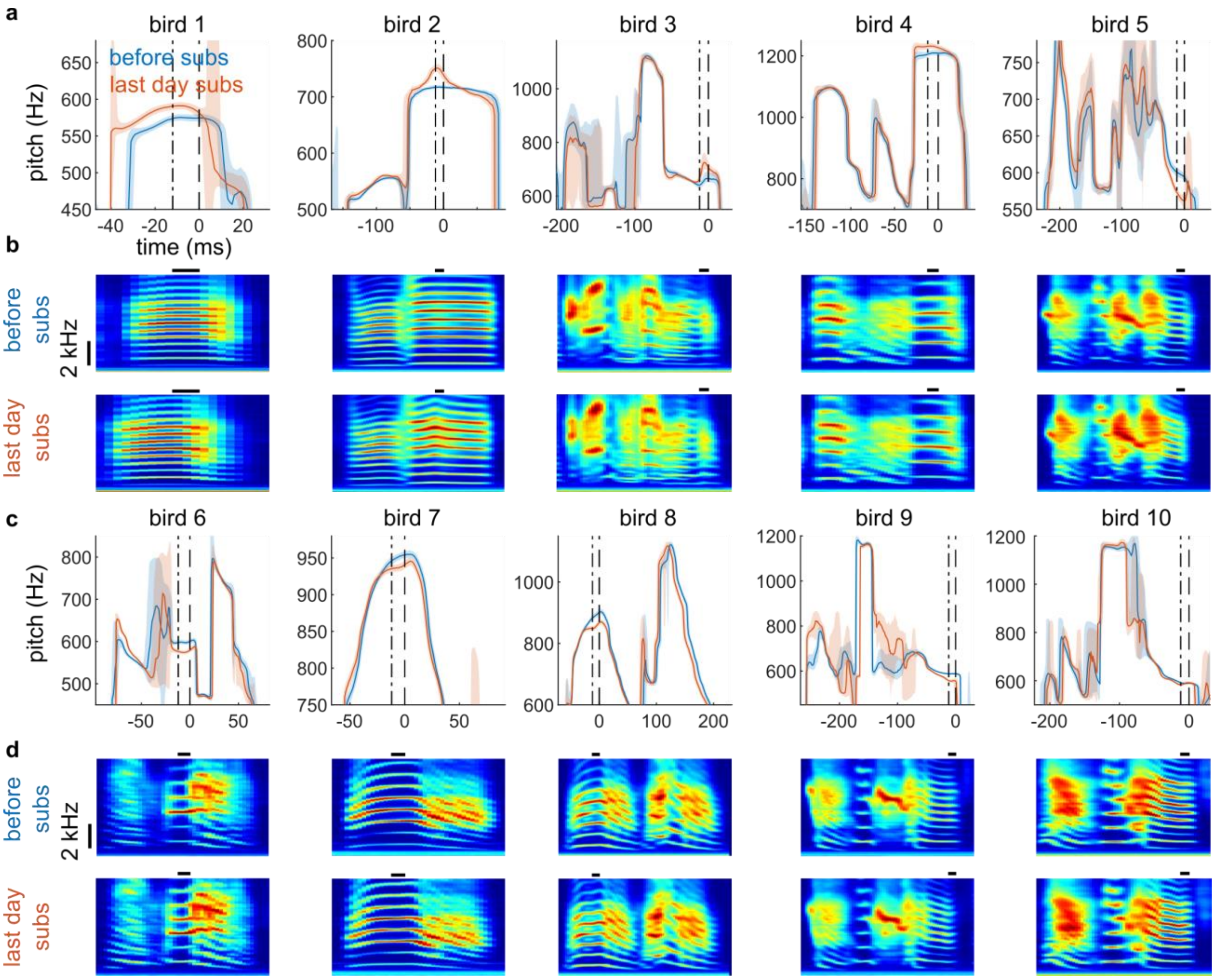


***Fig. S2: Within-syllable pitch trajectories in n=10 subs birds. a** and **c**, median (solid line) and quantiles (shaded area) of within-syllable pitch before substitution (before subs, blue) and on the last day of substitution (last day subs, red) in deaf birds subjected to subs for high-pitched syllables (a) and low-pitched syllables (b). The two dashed vertical lines show the window within which the pitch was calculated to determine whether light was switched off. The birds are ordered according to the blue/green bars in Fig. 1g. **b** and **d**. Average spectrogram of target syllable before (top row) and after (bottom row) the light off paradigm. Same time axis as in panel a. Horizontal black line shows the window where pitch is calculated.*

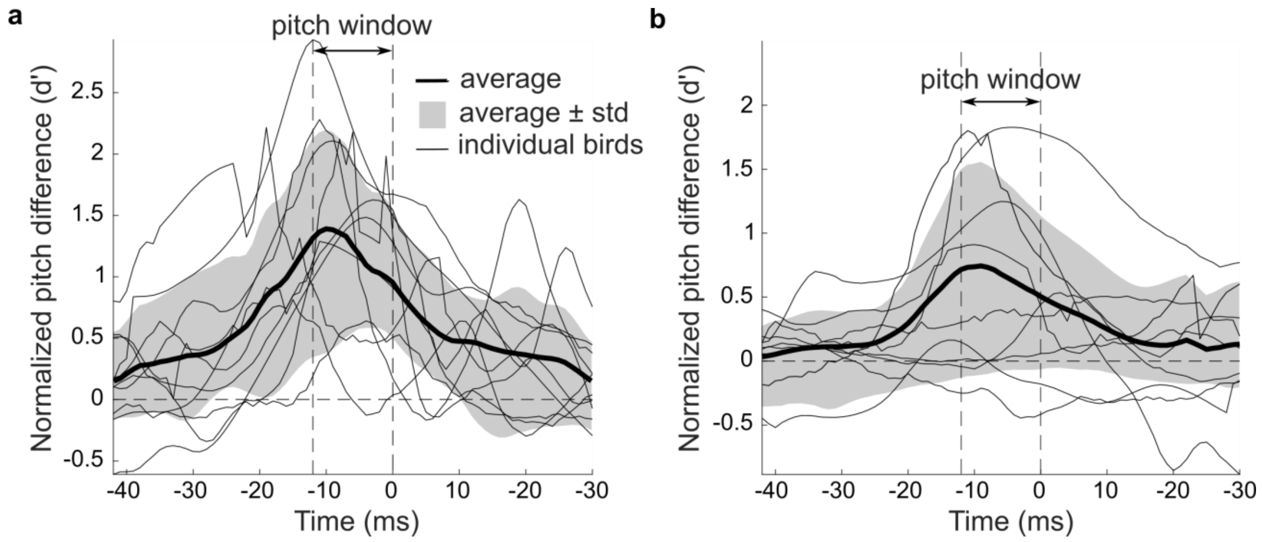


***Fig. S3: Traces in pitch difference show time-localized learning. a.*** *In subs birds, the difference trace in normalized pitch before light off and on the last day of light off reveals that birds adapt pitch within about 10 ms of the time window targeted for light off (dashed vertical lines). The fine curves show pitch difference traces in individual birds, the thick curve their average, and the gray area indicates ± one standard deviation. Curves in birds that decreased pitch are flipped to make differences positive.* ***b****. Same as in a but in hearing birds.*
